# Increasing glucose uptake into neurons antagonizes brain aging and promotes health and life span under dietary restriction in *Drosophila*

**DOI:** 10.1101/2020.06.11.145227

**Authors:** Mikiko Oka, Emiko Suzuki, Akiko Asada, Taro Saito, Koichi M. Iijima, Kanae Ando

**Affiliations:** Department of Biological Sciences, Graduate School of Science, Tokyo Metropolitan University, Tokyo, Japan; Department of Biological Sciences, Faculty of Science, Tokyo Metropolitan University, Tokyo, Japan; Gene Network Laboratory, National Institute of Genetics, Mishima, Shizuoka, Japan; Department of Genetics, SOKENDAI, Mishima, Shizuoka, Japan; Department of Alzheimer’s Disease Research, National Center for Geriatrics and Gerontology, Obu, Aichi, Japan; Department of Experimental Gerontology, Graduate School of Pharmaceutical Sciences, Nagoya City University, Nagoya, Aichi, Japan

**Keywords:** aging, brain, ATP, glucose transporter, glycolysis, mitochondria, dietary restriction, *Drosophila*

## Abstract

Brain neurons play a central role in organismal aging, but there is conflicting evidence about the roles of neuronal glucose availability, glucose uptake and metabolism in aging and extended lifespan. Here, we analyzed metabolic changes in the brain neurons of *Drosophila* during aging. Using a genetically-encoded fluorescent ATP biosensor, we found decreased ATP concentration in the neuronal somata of aged flies, which was correlated with decreased glucose content, expression of glucose transporter and glycolytic enzymes and mitochondrial quality. The age-associated reduction in ATP concentration did not occur in brain neurons with suppressed glycolysis or enhanced glucose uptake, suggesting these pathways contribute to reductions in ATP. Despite age-associated mitochondrial damage, increasing glucose uptake maintained ATP levels, suppressed age-dependent locomotor deficits and extended the life span. Increasing neuronal glucose uptake during dietary restriction resulted in the longest lifespans, suggesting an additive effect of enhancing glucose availability during a bioenergetic challenge on aging.

## Introduction

Aging is associated with progressive declines in physiological integrity and functions alongside increases in vulnerability and the risk of developing a number of diseases (Kirkwood and Austad, 2000). The nervous system regulates motor and sensory functions and orchestrates metabolic control and stress responses in peripheral tissues *via* neuroendocrine signals (Alcedo and Kenyon, 2004; Burkewitz et al., 2015; Cohen et al., 2009; Libert et al., 2007; Satoh and Imai, 2014; Ulgherait et al., 2014; Zhang et al., 2018). Thus, age-associated changes in neurons are likely to play a critical role in organismal aging.

The brain requires a large amount of energy: it consumes about 20% of the oxygen and 25% of the glucose in the human body (Belanger et al., 2011; Mink et al., 1981; Raichle et al., 1970). Most of the ATP required to support neuronal function is supplied by the metabolism of glucose through glycolysis and mitochondrial oxidative phosphorylation (Belanger et al., 2011). However, aging induces changes in both glucose availability and energy production, including declines in glucose uptake, electron transport chain activity, and aerobic glycolysis in the brain (Goyal et al., 2017; Hoyer, 2000). This implies that strategies aimed at increasing glucose metabolism in neurons may protect against organismal aging.

By contrast, dietary restriction (DR) and a reduction in insulin signaling have been demonstrated to have anti-aging effects (Guarente, 2008). DR causes circulating glucose concentrations to fall (Guarente, 2008). Thus, the pro-aging effects of reductions in brain glucose metabolism and the anti-aging effects of reducing insulin-stimulated glucose uptake are apparently contradictory. To resolve this discrepancy, it is necessary to elucidate how aging affects glucose metabolism in brain neurons and how such age-related changes in neurons interact with the anti-aging effects of DR.

In the present study, we show that aged brain neurons suffer from ATP deficits. Using a genetically encoded fluorescent ATP biosensor, we show that ATP concentrations decrease in the somas of brain neurons during aging due to a reduction in glycolysis or glucose availability. Increased neuronal glucose uptake by expression of a glucose transporter *hGlut3* is sufficient to ameliorate age-dependent declines in ATP via glycolysis. Of particular interest, we demonstrate that increasing glucose uptake into neurons significantly extends the organismal lifespan under DR conditions, suggesting that increasing neuronal glucose metabolism optimizes the anti-aging effects of energetic challenges.

## Results

### An ATP biosensor (ATeam) identifies local changes in ATP levels in Drosophila brain neurons

Neurons are highly polarized structurally and functionally, and require tight local management of ATP production (Rangaraju et al., 2014). To determine the local ATP concentration in fly brain neurons, we used the fluorescence resonance energy transfer (FRET)-based ATP biosensor, ATeam (Tsuyama et al., 2013). ATeam detects changes in ATP concentrations in the range of 0.5 to 4 mM at 25°C (Tsuyama et al., 2013; Tsuyama et al., 2017), which is appropriate for the physiological ATP concentrations in neurons (Erecinska and Silver, 1989; Pathak et al., 2015; Rangaraju et al., 2014). The concentrations of intracellular ATP are much higher than the expression levels of ATeam; therefore, the FRET signal derived from ATeam is not likely to be affected by its intracellular distribution (Tsuyama et al., 2013; Tsuyama et al., 2017).

It has been reported that the ATeam signal decreases in response to the inhibition of cellular respiration in the neurons of *Drosophila* larvae (Tsuyama et al., 2013; Tsuyama et al., 2017). To confirm that ATeam is capable of detecting changes in ATP concentration in adult fly brains, we used the Gal4-UAS system to express ATeam in neurons under the control of a pan-neuronal driver elav-GAL4 and inhibited respiratory complex III by treatment with antimycin (AM). We focused on the mushroom body, where axons, dendrites, and cell bodies are easily identifiable as lobes, calyxes, and Kenyon cells, respectively. AM treatment significantly reduced the FRET signal from ATeam in both the cell bodies and axons (Figure 1A), which indicates that ATeam can identify reductions in ATP concentrations in fly brain neurons.

**Figure 1.**
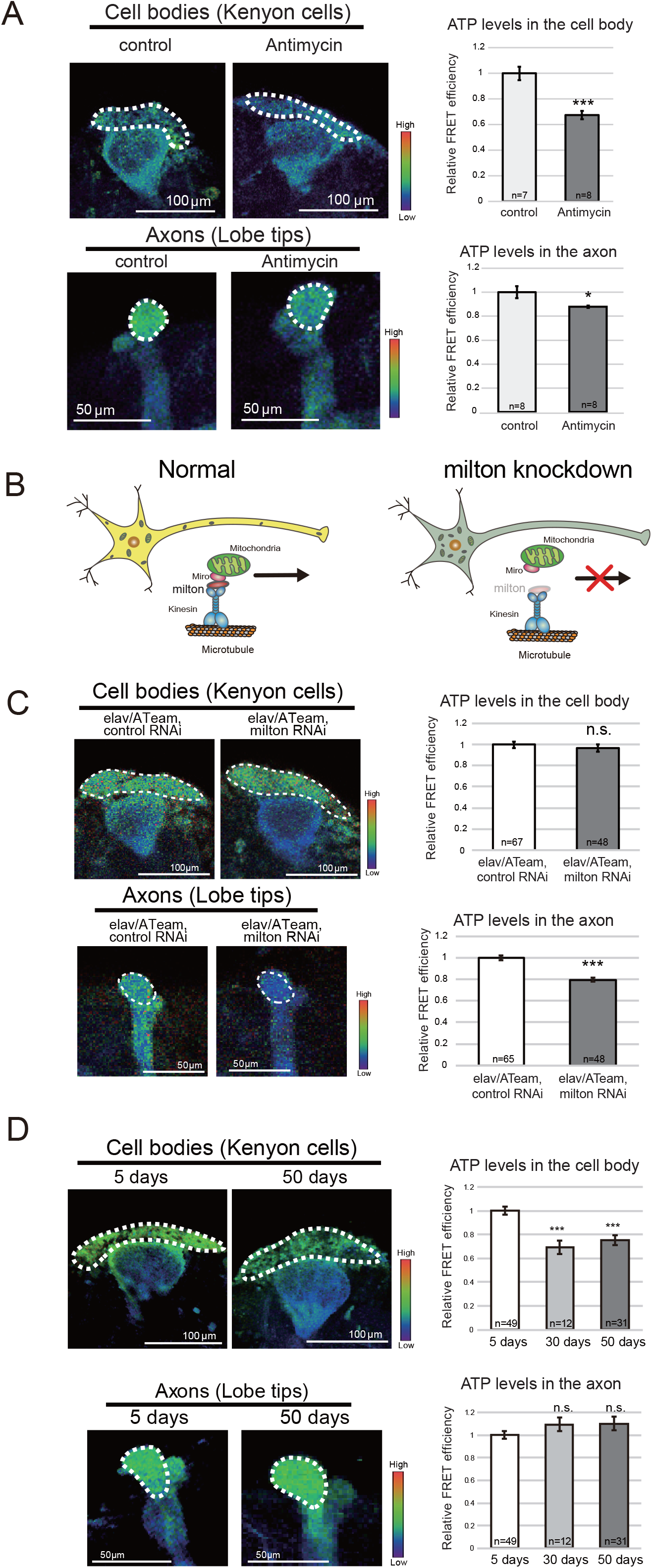
ATP concentration decreases in the cell body but not in the axon during aging. (A) A reduction in ATP concentration, caused by the inhibition of mitochondrial respiration by antimycin in adult fly brain neurons, was identified using ATeam. ATeam expression was driven by the pan-neuronal driver elav-GAL4. Quantification of the FRET signal (right panels) is shown (mean ± SE, n=7–8; *p<0.05, ***p<0.001; Student’s *t*-test). (B) Schematic representation of mitochondrial transport. The knockdown of milton, an adaptor protein for mitochondrial transport, depleted mitochondria in the axon. (C) Neuronal knockdown of milton reduced ATP concentration in Kenyon cells and lobe tips. Flies carrying the driver with control RNAi (elav/ATeam, control RNAi) were used as controls. Data are mean ± SE, n=48–65; n.s., p>0.05, ***p<0.001; Student’s *t*-test). (D) The FRET efficiency of ATeam in Kenyon cells (A) and lobe tips (B) of the mushroom body of 5-, 30-, and 50-day-old flies (n=12–49; n.s., p>0.05, ***p<0.001; one-way ANOVA, followed by Tukey’s HSD multiple comparisons test). ATeam expression was driven in neurons by the pan-neuronal driver Elav-GAL4.

Next, we determined whether ATeam could identify local changes in ATP concentration. To reduce ATP concentration in the axons only, axonal mitochondria were genetically depleted by knocking down milton, an adaptor protein for the kinesin motor that is essential for mitochondrial anterograde transport (Figure 1B) (Stowers et al., 2002). Previously, RNAi-mediated neuronal knockdown of milton was reported to effectively deplete mitochondria in axons (Iijima-Ando et al., 2012). Likewise, we found the FRET signal from ATeam decreased in axons, but not in the cell bodies, of milton-deficient fly brain neurons (Figure 1C). These results indicate that ATeam is capable of identifying local changes in ATP concentration in adult brain neurons.

### ATP concentration decreases in the cell body, but not in the axon, of brain neurons during aging

To determine whether ATP concentration changes in brain neurons in an age-related manner, we compared the ATP concentrations in the mushroom body structure of young (5-day-old), middle-aged (30-day-old), and old (50-day-old) flies. We found that the ATP concentration in the cell bodies in the mushroom body structure of middle-aged (30-day-old) and old (50-day-old) flies was ~70% of that in young flies (5-day-old) (Figure 1D). By contrast, there were no significant changes in axonal ATP concentrations during aging (Figure 1D). ATeam detects ATP levels as CFP and YFP signal ratio, suggesting that the reduction in ATeam signal was not due to the loss of neurons. To confirm this, we carried out immunostaining of the neuronal marker elav and ultrastructural analyzes of mushroom body region with electron microscopy. There were no vacuoles or signs of neuronal loss (Figure S1), indicating that the brains of 50-day-old flies were structurally intact, as previously reported (He and Jasper, 2014; Tonoki and Davis, 2015; White et al., 2010). Taken together, these results show that the ATP concentration, specifically in the cell bodies of brain neurons, decreases during aging.

### Decreases in glycolytic gene expression contribute to the age-related reduction in ATP

Most of the ATP produced in the brain is supplied by the metabolism of glucose (Belanger et al., 2011). To gain insight into the underlying reduced ATP concentrations, we measured glucose availability in the brain decreases during aging in *Drosophila*. We found that the glucose concentration in head extracts was significantly lower in aged flies (30- 50-day-old) than in young (5-day-old) flies (Figure 2A). Glucose uptake by brain neurons is mostly constitutive because the glucose transporter isoforms expressed in CNS neurons, Glut1 and Glut3 in mammals and Glut1 in *Drosophila*, are constitutively localized to the plasma membrane and insensitive to insulin (Chintapalli et al., 2007; Flier et al., 1987). We measured *dGlut1* mRNA expression during aging using qRT-PCR, which revealed that its expression was slightly but significantly lower in the heads of aged than in those of young flies (Figure 2B).

**Figure 2.**
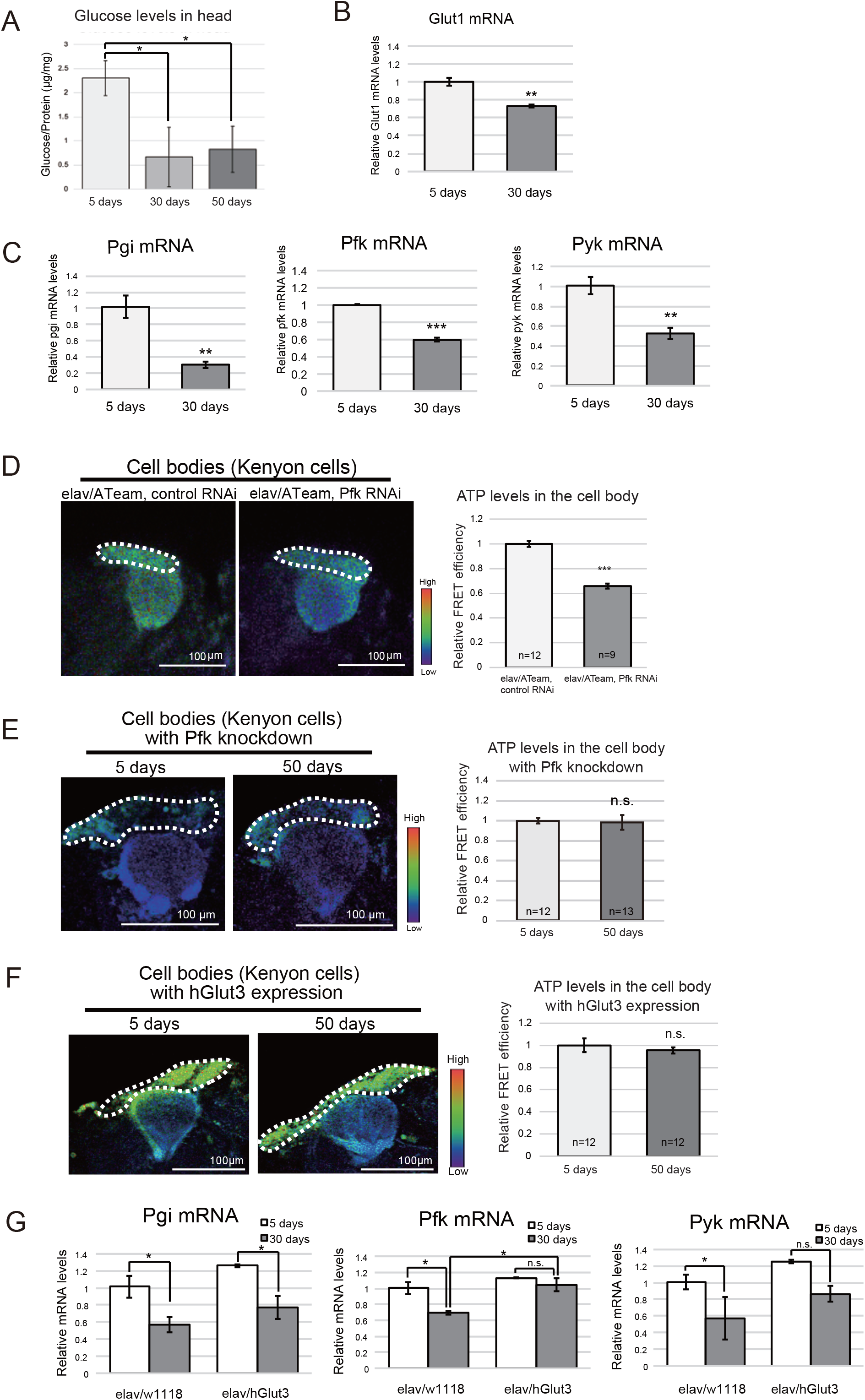
Neuronal expression of the glucose transporter *hGlut3* attenuates the age-related reduction in ATP concentration in brain neuron and glycolytic gene expression. (A) Glucose concentration in head lysates decreased during aging. The glucose concentrations in 5-, 30- 50-day-old fly heads, normalized to protein concentration, are shown (mean ± SE, n=6; *p<0.05; one-way ANOVA, followed by Tukey’s HSD multiple comparisons test). (B) *dGlut1* mRNA expression decreased in fly heads during aging. qRT-PCR was performed using RNA extracted from the heads of 5- 30-day-old flies (mean ± SE, n=3; **p<0.01; Student’s *t*-test). (C) The expression of glycolytic genes decreased in the fly brain during aging. The mRNA expression of genes encoding glycolytic enzymes in the heads of 5- 30-day-old flies was quantified using qRT-PCR (mean ± SE, n=3; **p<0.01, ***p<0.001; Student’s *t*-test). (D) Neuronal knockdown of *Pfk*, which encodes a rate-limiting glycolytic enzyme, reduced ATP concentration in 3-day-old fly neurons (mean ± SE, n=9–12; ***p<0.001; Student’s *t*-test). (E) The age-related reduction in ATP concentration in neuronal cell bodies was not identified in *Pfk*-deficient neurons. The expression of ATeam and *Pfk* RNAi was driven by the pan-neuronal driver elav-GAL4. The 5-day-old and 50-day-old flies were subjected to imaging analyzes at the same time. The experiments were repeated with timely independent cohorts of flies obtained from different crosses to analyze more than 12 brains for each age. Data are mean ± SE; n.s., p>0.05, *p<0.05; Student’s *t*-test. (F-G) Neuronal expression of *hGlut3* ameliorates the age-related reduction in ATP concentration and glycolytic gene expression and the decline in physical function. (F) ATP concentration did not decrease with aging in neurons overexpressing *hGlut3*. Expression of ATeam and hGlut3 was driven by the pan-neuronal driver elav-GAL4. The 5-day-old and 50-day-old flies were subjected to imaging analyzes at the same time. The experiments were repeated with timely independent cohorts of flies obtained from different crosses to analyze more than 12 brains for each age. (mean ± SE; n.s., p>0.05; Student’s *t*-test). (G) Neuronal expression of *hGlut3* attenuated the age-associated decline in glycolytic gene expression. The mRNA expression of genes encoding key glycolytic enzymes was quantified using qRT-PCR in the heads of 5- 30-day-old flies. Flies carrying the driver alone were used as controls. Data are mean ± SE, n=3; n.s., p>0.05, *p<0.05, **p<0.01; one-way ANOVA, followed by Tukey’s HSD multiple comparisons test.

In human brains, the rate of glycolysis changes during aging (Goyal et al., 2017). To determine the contribution of changes in glycolytic rate to the age-related reduction in ATP in the cell bodies of *Drosophila* brain neurons, we measured the expression of genes encoding the key glycolytic enzymes in young and aged fly heads including glucose-6-phosphate isomerase (*Pgi*), phosphofructokinase (*Pfk*), and pyruvate kinase (*Pyk*). qRT-PCR analysis revealed that the mRNA expression of *Pfk* was significantly lower in aged flies than in young flies (Figure 2C). To evaluate the contribution of glycolysis to ATP production in brain neurons, we knocked down *Pfk*, which encodes phosphofructokinase, a rate-limiting enzyme in glycolysis. Neuronal knockdown of *Pfk* reduced ATP concentration by up to 65% in 3-day-old flies (Figure 2D), indicating that glycolysis plays a critical role in ATP production in brain neurons. To determine whether the decline in glycolytic activity contributes to the age-related decrease in ATP concentration, we measured the ATeam signal in *Pfk*-deficient young and aged brain neurons. The neurons of 5-day-old flies with *Pfk* knockdown showed a reduced ATP concentration, with no further reduction in 50-day flies (Figure 2E). This result suggests that a lower rate of glycolysis and/or events upstream of glycolysis are responsible for the age-related reduction in ATP in cell bodies of brain neurons.

### Neuronal expression of the glucose transporter hGlut3 attenuates the age-related reduction in ATP concentration in brain neurons and glycolytic gene expression

We next asked whether increasing glucose uptake could ameliorate the age-related reduction in ATP in the brain neurons. Expression of *hGlut3* in *Drosophila* cells has been reported to increase glucose uptake (Besson et al., 2015). We found that neuronal rexpression of *hGlut3* ameliorated the age-related reduction in ATP in the neuronal cell bodies (Figure 2F). By contrast, neuronal oexpression of *hGlut3* did not increase the ATP concentration in young flies (Figure S2). These results suggest that glucose availability is the rate-limiting factor specifically in the brain neurons of old flies, and that overexpression of glucose transporter can increase ATP production in the neurons of aged flies.

We also examined whether glucose transporter overexpression attenuates the age-related changes in glycolytic activity. We found that neuronal expression of *hGlut3* suppressed the age-related decrease in the mRNA expression of the glycolytic genes, *Pgi*, *Pfk*, and *Pyk* (Figure 2G). Taken together, these results suggest that neuronal overexpression of *hGlut3* ameliorates age-related declines in ATP concentration by increasing glucose metabolism in aged fly neurons.

### The age-related increase in abnormal mitochondria is not reversed by overexpression of glucose transporter in neurons

Since mitochondrial dysfunction is implicated in aged brains, we examined whether the reduced number and increased damage to mitochondrial was associated with an age-related reduction in ATP levels in brain neurons. We first measured the mitochondrial density in the brain neurons of young and aged flies by transmission electron microscopy (TEM). In the cell body, where ATP concentrations decreases (Figure 1D), the number of mitochondria did not change significantly during aging (Figure 3A). Thus, the age-related reduction in ATP in the cell bodies of neurons was not the result of a decrease in the number of mitochondria.

**Figure 3.**
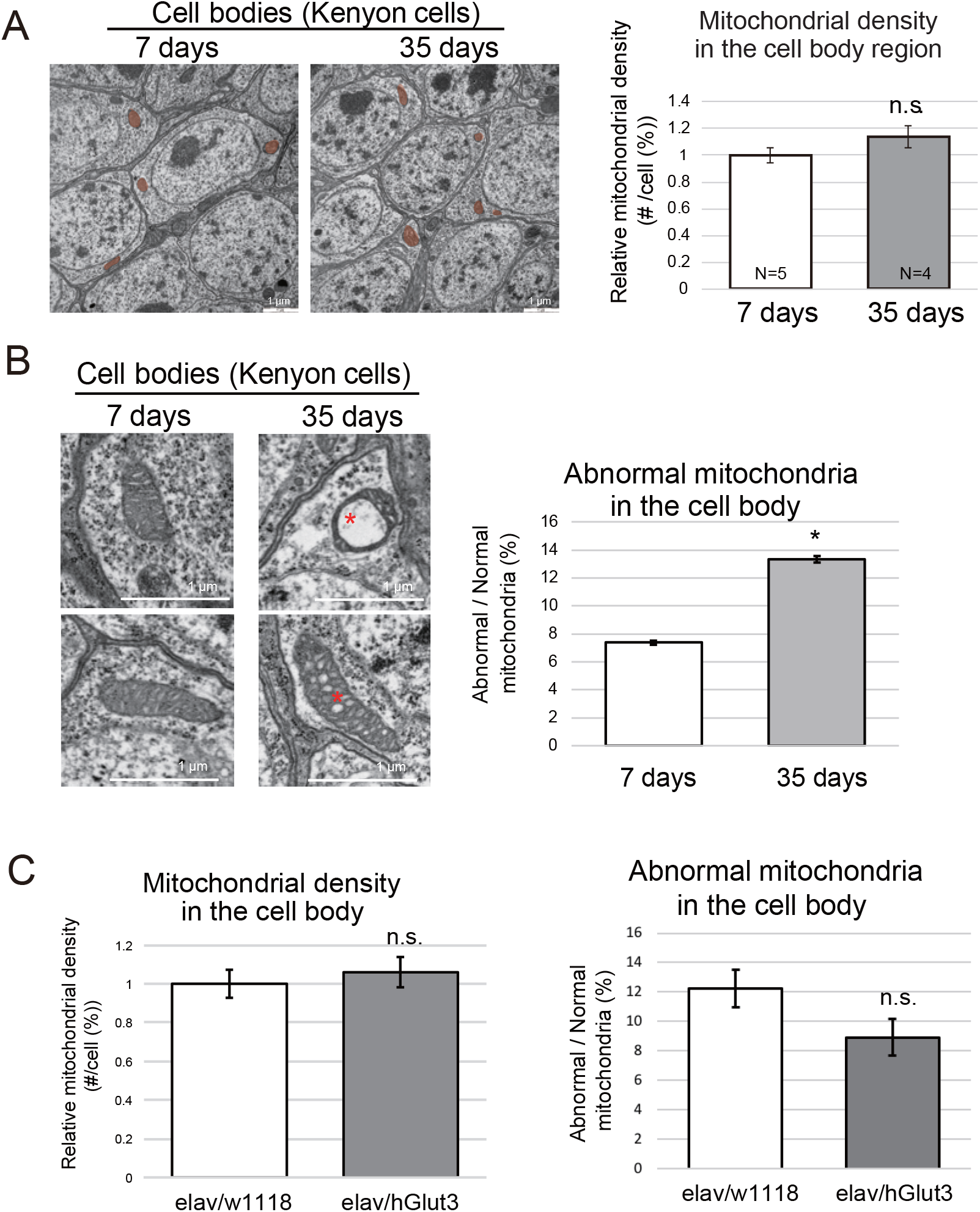
Age-related changes in the number and quality of mitochondria in brain neurons. (A) The number of mitochondria in the neuronal cell body did not significantly change during aging. More than four brain hemispheres from flies of each age were analyzed using transmission electron microscopy (TEM), and the numbers of mitochondria in more than 30 cells in each hemisphere were counted. Representative images are shown, with the mitochondria highlighted in orange. (B) The ratio of the number of abnormal mitochondria to the total number of mitochondria increased in the cell body during aging. More than four brain hemispheres from flies of each age were examined using TEM, and more than 30 cells were analyzed in each hemisphere. The number of mitochondria containing vacuoles or disorganized cristae (asterisks) was counted. (mean ± SE, n=4–5; n.s., p>0.05, *p<0.05; Student’s t-test). (C) Neuronal overexpression of *hGlut3* did not affect the age-associated changes in mitochondria in the brain. The mitochondrial density (left) and the ratio of the number of abnormal mitochondria to the total number of mitochondria (right) in neurons expressing *hGlut3* in the Kenyon cell region of the brains were analyzed under TEM 50 days after eclosion (mean ± SE, n=4–6; n.s., p>0.05; Student’s t-test).

Next, we analyzed the quality of mitochondria in the brain neurons of young and aged flies. Dysfunctional mitochondria are recognized by the abnormal structure of their cristae (Brandt et al., 2017; Daum et al., 2013). Ultrastructural analyzes of the mitochondria revealed that the number of abnormal mitochondria in cell bodies slightly but significantly increased during aging (7% in 7-day-old flies and 13% in 35-day-old flies, Figure 3B). By contrast, in the axon, where ATP concentration does not decrease (Figure 1D), the number of mitochondria decreased slightly, but the number of abnormal mitochondria did not significantly change during aging (Figure S3), which might be because of the presence of a robust quality control system in this location (Lin et al., 2017). These results indicate that a slight but significant increase in the number of damaged mitochondria occurs alongside a decrease in ATP concentration in the cell bodies of neurons during aging.

Interestingly, while neuronal overexpression of *hGlut3* ameliorated the age-related reduction in ATP in the neuronal cell bodies (Figure 2F), it neither increased the number of total mitochondria, nor reduced the ratio of abnormal mitochondria (Figure 3C). These results suggest that, increasing glucose uptake is sufficient to ameliorate energetic deficits in aged brain neurons.

### Increasing neuronal glucose uptake ameliorates declines in physical performance in aged flies and has an additive effect with DR in extending lifespan

Finally, we examined whether the increase in glucose uptake into neurons promoted the health and life span of flies. We first analyzed the effects of neuronal overexpression of *hGlut3* on the age-related decline in physical function using a climbing assay, a robust behavioral assay taking advantage of the innate negative geotaxis behavior. In wild-type flies, climbing ability declined by 45 days of age, whereas flies with neuronal expression of *hGlut3* performed significantly better at this age (Figure 4A), indicating that the increase in glucose uptake into neurons promoted healthspan in flies. We also found that flies with neuronal expression of *hGlut3* lived slightly but significantly longer than control flies (Figure 4B).

**Figure 4.**
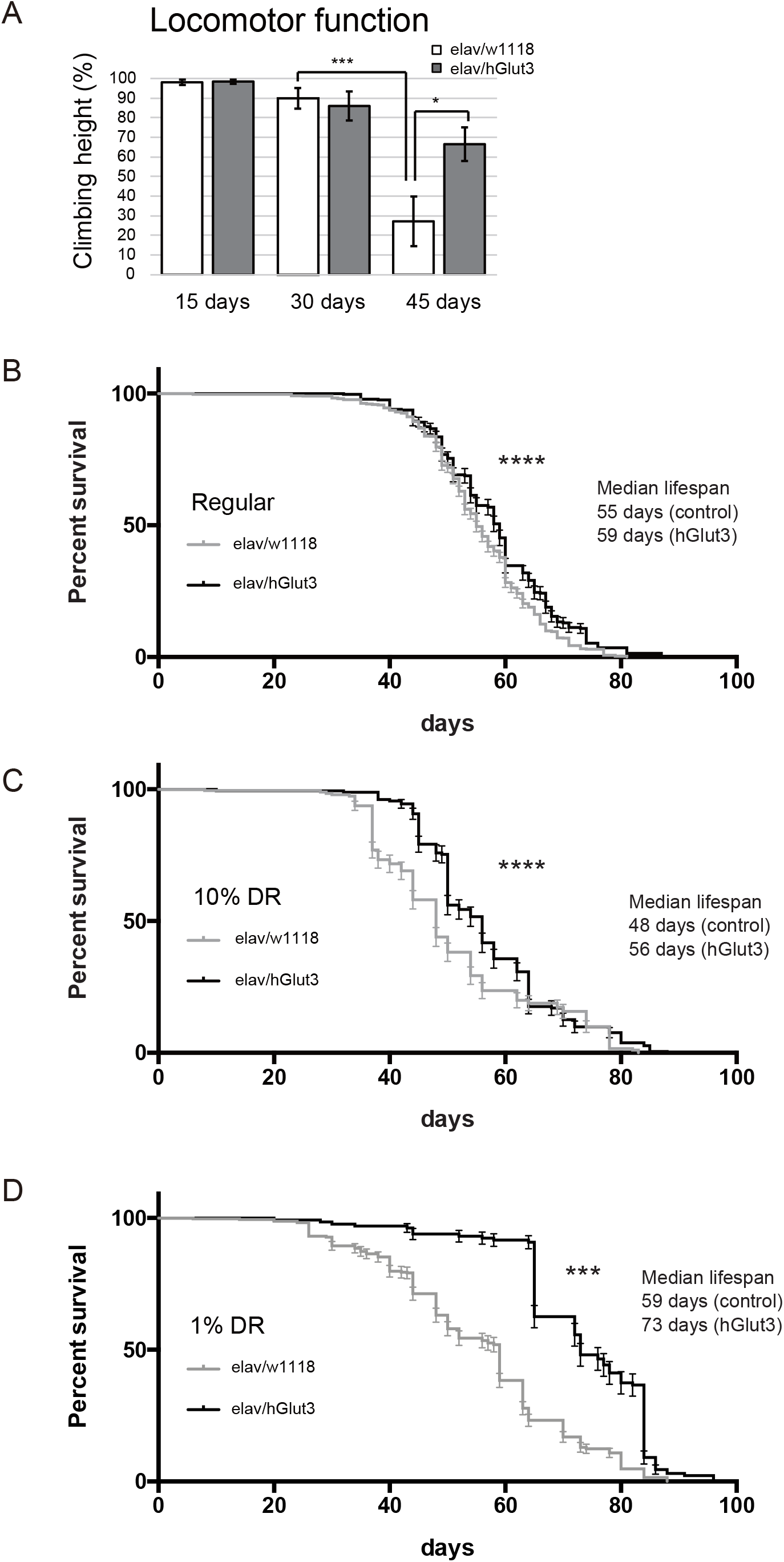
Increasing neuronal glucose uptake attenuates declines in physical performance in aged flies and has an additive effect with dietary restriction (DR) in extending lifespan. (A) Neuronal expression of *hGlut3* ameliorated the age-associated neuronal dysfunction. Flies were tapped to the bottom of the vial, and the distance that they then climbed in 30 seconds was measured (mean ± SE, n=12–39; *p<0.05, ***p<0.001; Student’s *t*-test). Flies carrying the driver alone were used as controls. (B-D) Neuronal expression of *hGlut3* and DR synergistically extended lifespan. Flies were raised on regular cornmeal food and maintained on regular cornmeal food (B) or two types of DR diet (C and D) (C: food without cornmeal and containing 10% (w/v) of yeast and glucose (10%), D: food without cornmeal and containing 1% (w/v) of yeast and glucose (1%) after eclosion. Error bars represent 95% confidence intervals. Data are expressed as mean ± SE, n=131-601, ***; p<0.001, ****; p<0.0001, Log-rank test.

DR has been reported to extend the life span in most species, including *Drosophila* (Bauer et al., 2005). Since DR causes circulating glucose concentrations to fall (Guarente, 2008), the anti-aging effects of DR and increasing in glucose uptake into neurons might be independent and additive. To determine the relationship between neuronal uptake of glucose and DR in aging, we analyzed the effects of neuronal expression of *hGlut3* on life span under various dietary conditions. Flies were raised on regular cornmeal food, and after eclosion, they were fed regular food, DR food (10%) (10% yeast, 10% glucose, no cornmeal) or DR food (1%) (1% yeast, 1% glucose, no cornmeal). Flies on DR food showed reduced levels of glucose in the head and body with greater effects seen with 1% food than 10% food (Figure S4A). The life span was significantly extended on 1% food (Figure S4B) and life span extension by neuronal expression of *hGlut3* was more prominent under DR conditions. While the median lifespan on regular food was 55 days in control and 59 days in *hGlut3*-expressing flies maintained on regular food (Figure 4B), on 1% food, the median lifespan was 59 days in control and 73 days in *hGlut3*-expressing flies (Figure 4D). Maintaining flies on 10% food did not affect the life span of control flies, but it extended the life span of *hGlut3*-expressing flies (median lifespan 48 days in control and 56 days in *hGlut3*-expressing flies (Figure 4C)).

We also analyzed glucose levels after *hGlut3* expression in the aged flies under normal and DR conditions. We found that *hGlut3* expression in neurons did not restore glucose levels in aged flies under normal or DR conditions (Figure S5). This result suggests that lifespan extension caused by neuronal expression of *hGlut3* is not due to gross changes in total levels of glucose in the brain, but is associated with efficient uptake of glucose and its utilization in neurons. When glucose availability is limited under DR conditions, efficiency of glucose utilization would have more impact than under normal conditions. Enhancement of glucose uptake by overexpression of *hGlut3* may, therefore, be prominent under DR conditions. Taken together, these results suggest that increased glucose uptake by neurons enhances the anti-aging effects of DR at the organismal level.

## DISCUSSION

In the present study, we showed that aged brain neurons suffer from energy deficits, along with decreases in glucose concentration, the expression levels of glucose transporter and glycolytic enzymes (Figure 1–2) and mitochondrial quality (Figures 3). Increasing glucose uptake by neurons was sufficient to attenuate energetic deficits (Figure 2) despite persistent mitochondrial damage (Figures 3) and suppressed age-dependent locomotor decline (Figure 4). Interestingly, the effects of increasing glucose uptake on lifespan extension were more prominent under DR (Figure 4B-D), suggesting that increasing neuronal uptake of glucose in concert with bioenergetic challenges could benefit health and lifespan in flies.

During aging, the ATP supply to brain neurons decreases because of global reductions in glucose metabolism, including in oxygen consumption, glycolysis, and mitochondrial respiratory capacity (Conley et al., 2000; Goyal et al., 2017; Jang et al., 2018; Kety, 1956; Martin et al., 1991). Our results suggest that glucose uptake was a limiting factor for ATP supply in aged brain neurons. An increase in glucose uptake attenuated not only the reduction in neuronal ATP concentration during aging, but also the age-related reduction in the expression of glycolytic enzymes (Figure 2). In mammalian cells, it has been reported that an elevated concentration of intracellular glucose activates hypoxia-inducible factor-1α (Guo et al., 2008), which promotes the expression of glycolytic enzymes (Lum et al., 2007). An increase in glucose uptake did not suppress the age-related reduction in mitochondrial quality (Figure 3), suggesting that enhanced glucose supply can produce sufficient ATP despite mitochondrial damage to produce sufficient ATP *in vivo*.

DR is known to extend lifespan in many species, but it may reduce glucose supply to the brain and worsen energy deficits in neurons. We used DR conditions that are similar to Bauer et al. (Bauer et al., 2005), under which both the levels of sugar and yeast are reduced. Unlike Bauer et al., we saw an initial increase in mortality followed by a prolongation of lifespan (Figure S4), although the extension of lifespan was less than that observed in previous works (Bauer et al., 2005; Bross et al., 2005). This difference may be due to different laboratory media used to maintain stocks. The food used in our facility is relatively rich in yeast and cornmeal (4% yeast, 10% glucose, 9% cornmeal) compared with those used by Bauer et al. (2% yeast, 10% sucrose, 5% cornmeal); this difference may have affected the response to DR in the studied offspring.

Our results suggest that increased glucose uptake by neurons further improves the anti-aging effects of DR, presumably by counteracting neuronal energy deficits under DR conditions (Figure 4). Lower extracellular glucose caused by DR may recapitulate some of the conditions observed in the aged brain, in which lower cerebral blood flow limits glucose supply (Goyal et al., 2017; Hoyer, 2000; Noda et al., 2002). In *Drosophila*, lifespan extension by DR is mostly modulated by dietary yeast (Grandison et al., 2009; Lee et al., 2008; Mair et al., 2005): DR in *Drosophila* research is often performed simply by reducing the amount of yeast. In this study, both sugar and yeast were reduced in the DR condition, which caused a reduction in overall glucose availability (Figure S4). One might suspect that a reduction in sugar would have a negative effect on survival and *hGlut3* overexpression simply restored glucose availability. However, the glucose content in the head was not restored by neuronal expression of *hGlut3* (Figure S5), suggesting this was not the case. Instead, our results suggest that enhancing neuronal glucose uptake had more impact under conditions in which glucose availability is limited. In support of this, a previous study that did not use DR showed no effect of neuronal expression of *dGlut1* or *hGlut3* on lifespan (Besson et al., 2015; Niccoli et al., 2016).

The anti-aging effects of DR are reported to be mediated by a reduction in the intake of protein or amino acids (Mirzaei et al., 2014). Amino acid restriction affects lifespan through effects on amino-acid sensing pathways, such as mechanistic target of rapamycin complex 1 and the integrated stress response (Mirzaei et al., 2014). The additive effects of DR and neuronal expression of *hGlut3* on lifespan may be explained by the respective peripheral activation of anti-aging pathways by DR and increased neuronal glucose uptake and metabolism. Thus, a strategy aimed at both increasing glucose uptake by neurons and restricting amino acid intake may be effective at reducing the effects of aging.

In summary, we have shown that glucose uptake by brain neurons is a critical regulator of organismal and brain aging. Further elucidation of the cross-talk among nutrient signaling pathways in the central nervous system and peripheral tissues during aging will lead to the identification of novel strategies for the extension of the healthy lifespan of an organism.

## Acknowledgement

The authors thank Dr. Hiromi Imamura from the Graduate School of Biostudies, Kyoto University, for the ATP biosensor transgenic fly line and discussions. Authors thank Dr. Marie Thérèse Besson, Aix-Marseille Université, for UAS-hGlut3 fly line. Authors are grateful to Dr. Shin-ichi Hisanaga from the Department of Biological Sciences, Tokyo Metropolitan University, for critical comments. Authors thank Tetsuya Miyashita for technical support. This work was supported by a research award from the Japan Foundation for Aging and Health (to K.A) and a Grant-in-Aid for Scientific Research on Challenging Research (Exploratory) [JSPS KAKENHI Grant number 19K21593] (to K.A.), NIG-JOINT (71A2018, 25A2019) (to K.A.), and [JSPS Grant-in-Aid for JSPS Research Fellow 18J21936] to O.M.), and the Research Funding for Longevity Science 19-7 from the National Center for Geriatrics and Gerontology, Japan (to K.M.I.).

## Author Contributions

M.O., K.M.I and K.A. designed the experiments, interpreted the data, and wrote the manuscript; M.O., A.A., T.S. and E.S. conducted the experiments; and M.O., E.S. and K.A. analyzed the data. All authors read and approved the manuscript.

## Declaration of Interests

The authors declare no competing interests.

## Supplemental Information

**Figure S1. Kenyon cells in aged fly brains are structurally intact.** (A) Fly brains expressing an ATP biosensor in neurons were immunostained with an antibody for the neuronal marker elav. Low and high magnification of the maximum intensity projection of Calyx (CA) and Kenyon Cells (KC) are shown. More than six brains were analyzed, and no degeneration was observed. (B) Ultrastructural analyzes of Kenyon cells show no sign of cell death in the aged brains. More than four brain hemispheres from young (7-day-old) and aged (50-day-old) flies were analyzed using transmission electron microscopy (TEM).

**Figure S2. The overexpression of *hGlut3* does not increase ATP concentration in the mushroom body neurons of young flies.** The FRET signals from ATeam were quantified and are shown as mean ± SE (n=12, n.s.; p>0.05, Student’s *t*-test). The flies were 5 days old.

**Figure S3. Age-related changes in the number and quality of mitochondria in the axon.** (A) The number of mitochondria in the axon significantly decreased during aging. More than four brain hemispheres from flies of each age were analyzed using transmission electron microscopy (TEM), and the numbers of mitochondria in the lobe tip in at least in each hemisphere were counted. Representative images are shown, with the mitochondria highlighted in orange. (B) The ratio of the number of abnormal mitochondria to the total number of mitochondria did not significantly change in the axon during aging. The number of mitochondria containing vacuoles or disorganized cristae was counted. Scale bar: 1 μm. Mean ± SE, n=4–5; n.s., p>0.05, *p<0.05; Student’s *t*-test.

**Figure S4. The effects of dietary restriction (DR) on glucose levels and life span.** (A) Flies undergoing DR have lower levels of glucose in their heads. The glucose and protein concentrations in the heads of flies subjected to DR. Flies were raised on regular cornmeal food and after eclosion, were maintained on regular corn meal food (Regular) or two types of DR diet (10% and 1%)(10%: food without cornmeal and containing 10% (w/v) of yeast and glucose, 1%: food without cornmeal and containing 1% (w/v) of yeast and glucose). Flies were 30-day-old. Data are mean ± SE, n=3, **; p<0.01, ***; p<0.001, Student’s *t*-test. (B) Flies maintained on 1% food showed longer lifespan. Error bars represent 95% confidence intervals. Data are mean ± SE, n=131-601, n.s.; p>0.05, ****; p <0.0001, Log-rank test.

**Figure S5. Glucose levels in the head or body were not increased by neuronal expression of***hGlut3* **under regular or dietary restriction (DR) conditions.** Flies were raised on regular cornmeal food and maintained on regular cornmeal food or two types of DR diet (10% and 1%) (10%: food without cornmeal and containing 10% (w/v) of yeast and glucose, 1%: food without cornmeal and containing 1% (w/v) of yeast and glucose) after eclosion. The glucose concentrations in 30-day-old fly heads (A) and bodies (B) are shown (mean ± SE, n=3, Student’s t-test, p>0.05).

**Supplemental Table. The genotypes of flies used in each experiment.**

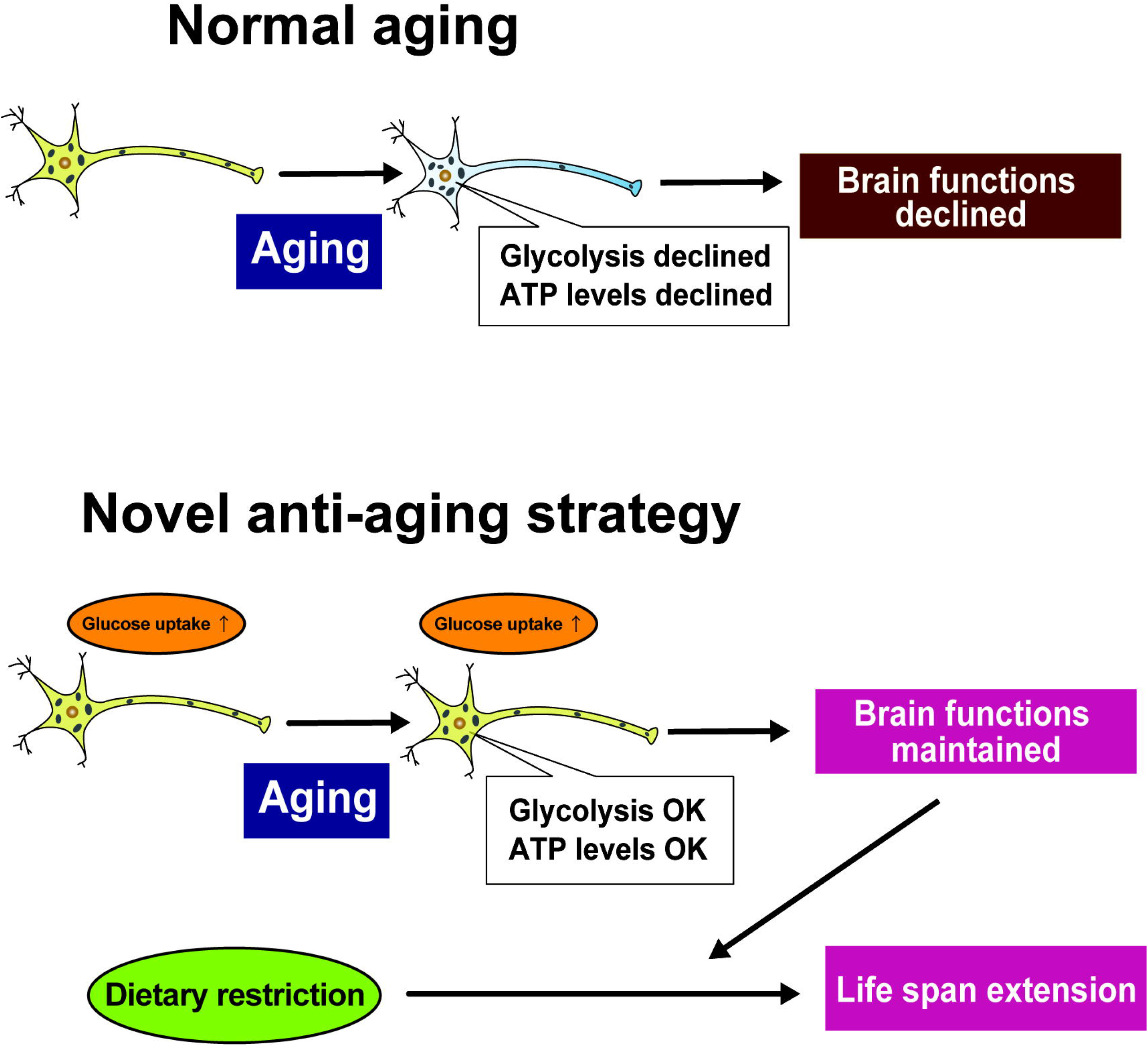

## Notes

### Competing Interest Statement

The authors have declared no competing interest.

