## Supplemental figures and table for "Increasing glucose uptake into neurons antagonizes brain aging and promotes health and life span under dietary restriction in *Drosophila*"

### Supplemental figure 1

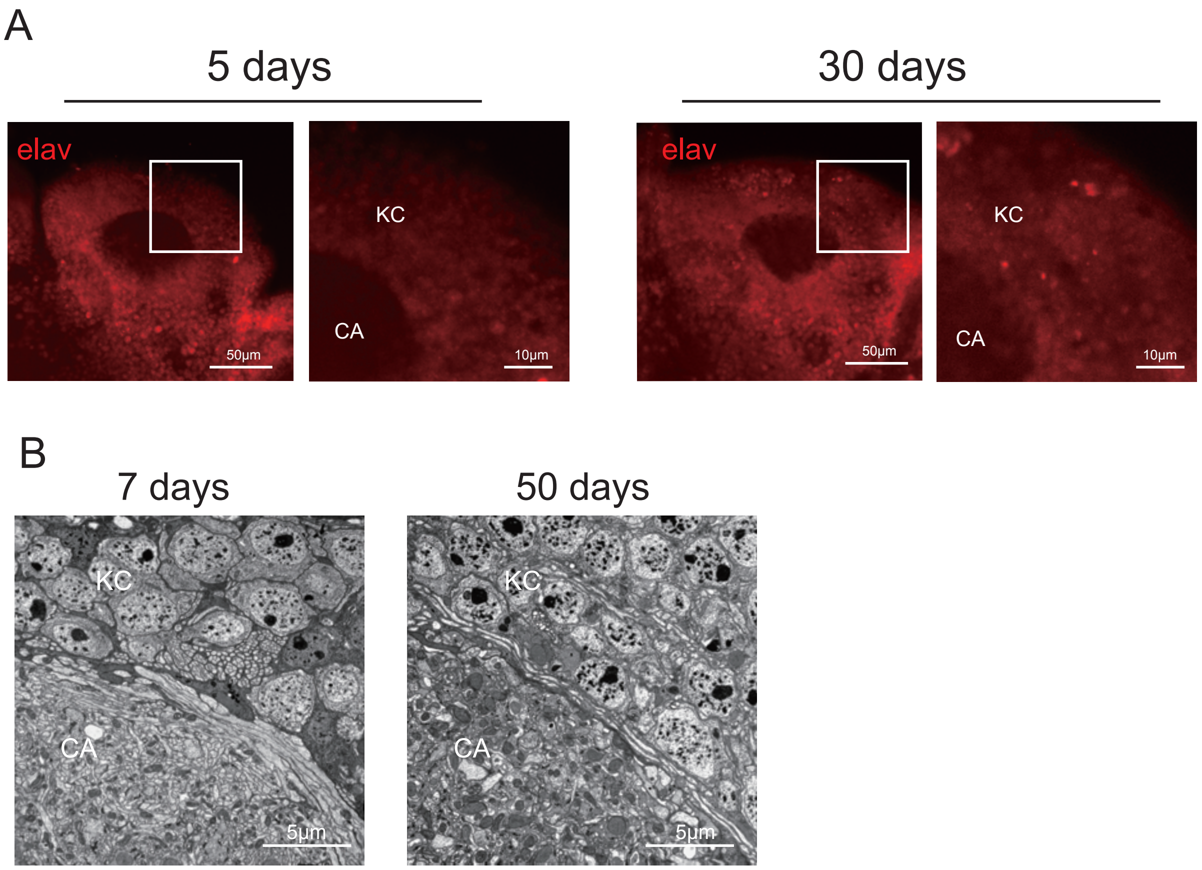


**Figure S1. Kenyon cells in aged fly brains are structurally intact.**
(A) Fly brains expressing an ATP biosensor in neurons were immunostained with an antibody for the neuronal marker elav. Low and high magnification of the maximum intensity projection of Calyx (CA) and Kenyon Cells (KC) are shown. More than six brains were analyzed, and no degeneration was observed. (B) Ultrastructural analyzes of Kenyon cells show no sign of cell death in the aged brains. More than four brain hemispheres from young (7-day-old) and aged (50-day-old) flies were analyzed using transmission electron microscopy (TEM).

### Supplemental figure 2

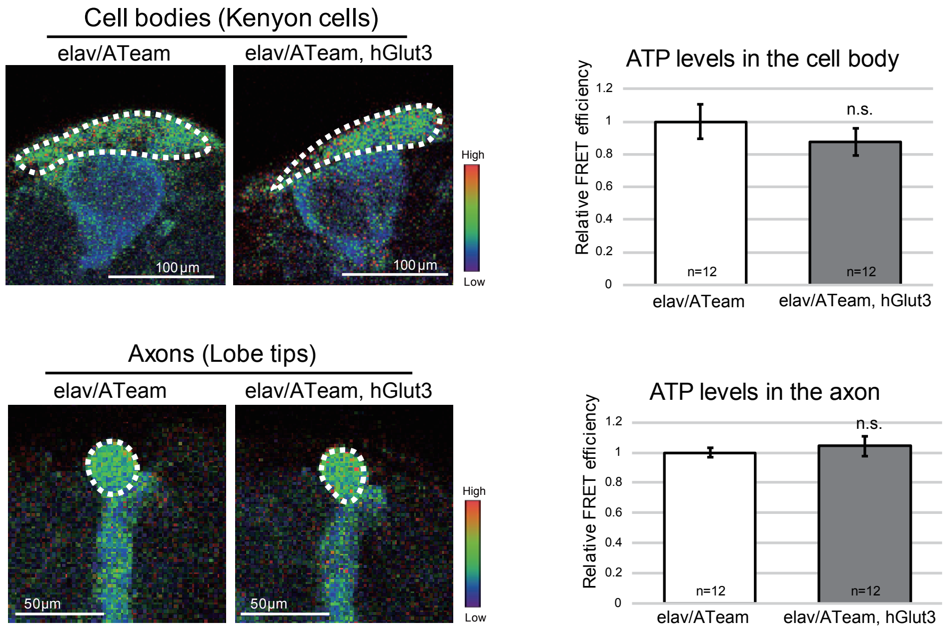


**Figure S2. The overexpression of hGlut3 does not increase ATP concentration in the mushroom body neurons of young flies.**
The FRET signals from ATeam were quantified and are shown as mean ± SE (n=12, n.s.; $p>0.05$, Student's t-test). The flies were 5 days old.

### Supplemental figure 3

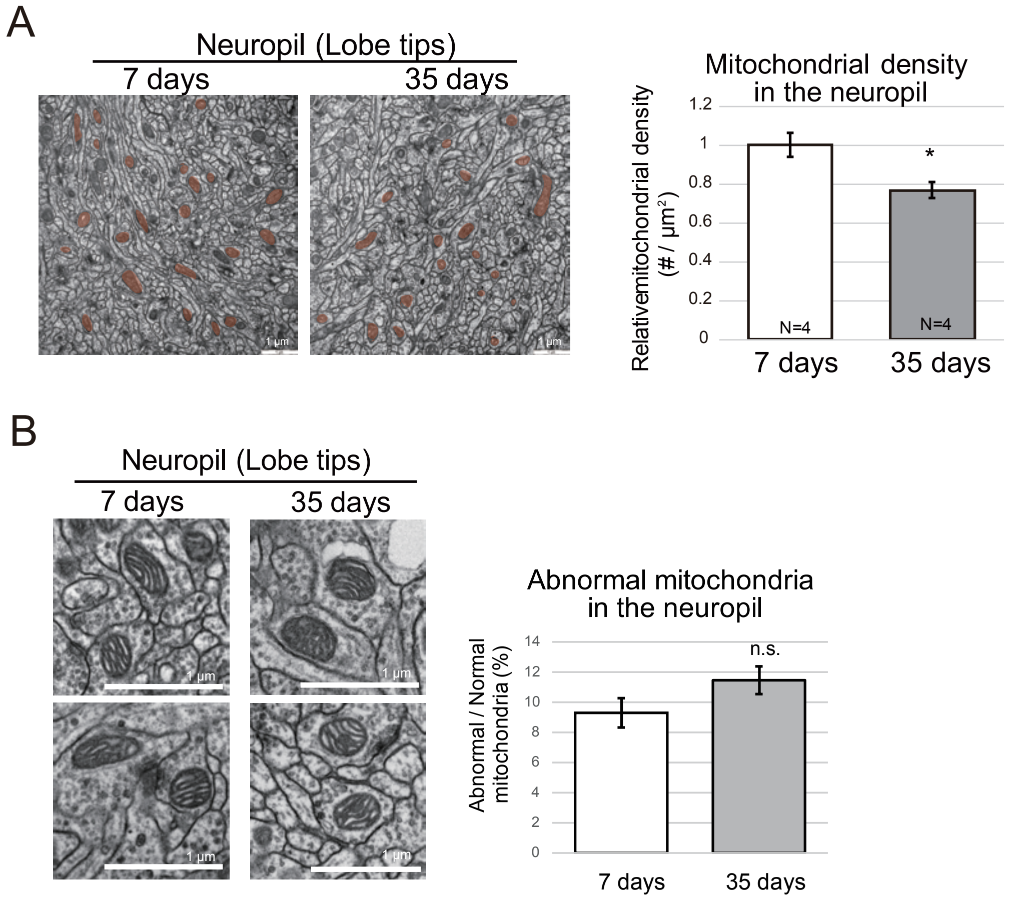


**Figure S3. Age-related changes in the number and quality of mitochondria in the axon.**

(A) The number of mitochondria in the axon significantly decreased during aging. More than four brain hemispheres from flies of each age were analyzed using transmission electron microscopy (TEM), and the numbers of mitochondria in the lobe tip in at least in each hemisphere were counted. Representative images are shown, with the mitochondria highlighted in orange.

(B) The ratio of the number of abnormal mitochondria to the total number of mitochondria did not significantly change in the axon during aging. The number of mitochondria containing vacuoles or disorganized cristae was counted. Scale bar: 1 μm. Mean ± SE, n=4–5; n.s., $p>0.05$, $^*p<0.05$; Student's t-test.

### Supplemental figure 4

A

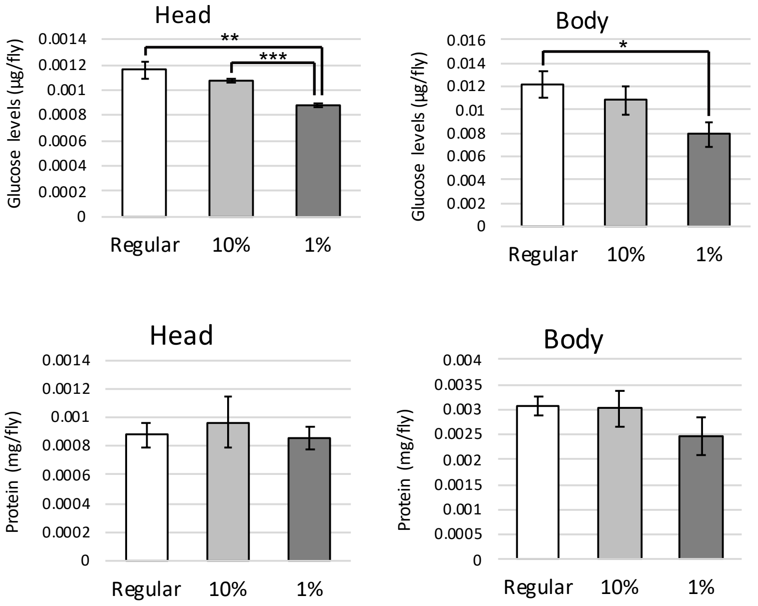


B

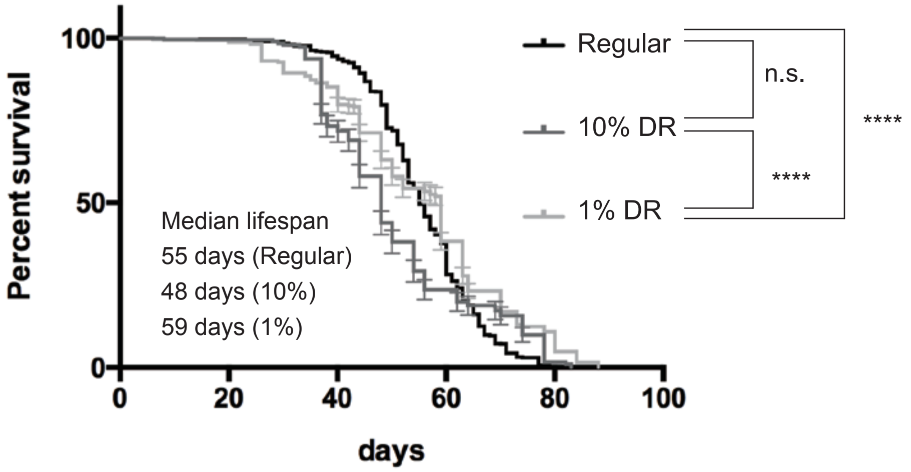


**Figure S4. The effects of dietary restriction (DR) on glucose levels and life span.** (A) Flies undergoing DR have lower levels of glucose in their heads. The glucose and protein concentrations in the heads of flies subjected to DR. Flies were raised on regular cornmeal food and after eclosion, were maintained on regular corn meal food (Regular) or two types of DR diet (10% and 1%)(10%: food without cornmeal and containing 10% (w/v) of yeast and glucose, 1%: food without cornmeal and containing 1% (w/v) of yeast and glucose). Flies were 30-day-old. Data are mean ± SE, n=3, **; p<0.01, ***; p<0.001, Student's t-test. (B) Flies maintained on 1% food showed longer lifespan. Error bars represent 95% confidence intervals. Data are mean ± SE, n=131-601, n.s.; p>0.05, ****; p <0.0001, Log-rank test.

### Supplemental figure 5

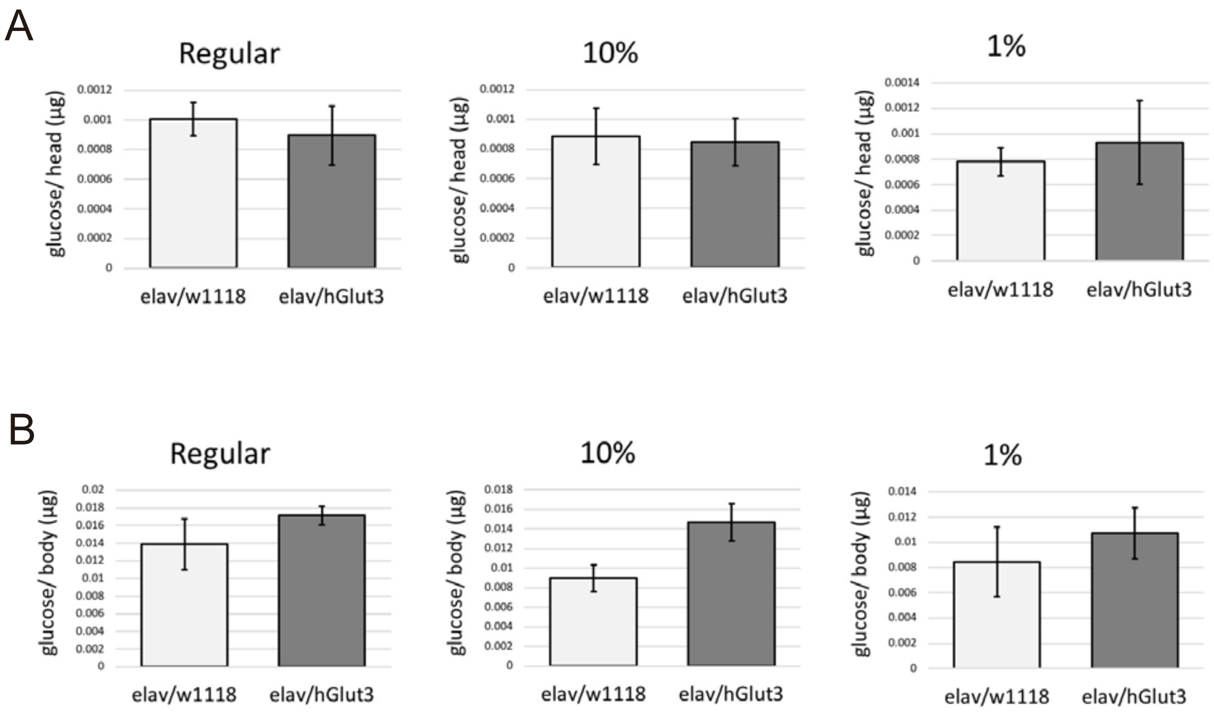


**Figure S5. Glucose levels in the head or body were not increased by neuronal expression of hGlut3 under regular or dietary restriction (DR) conditions.**

Flies were raised on regular cornmeal food and maintained on regular cornmeal food or two types of DR diet (10% and 1%) (10%: food without cornmeal and containing 10% (w/v) of yeast and glucose, 1%: food without cornmeal and containing 1% (w/v) of yeast and glucose) after eclosion. The glucose concentrations in 30-day-old fly heads (A) and bodies (B) are shown (mean ± SE, n=3, Student's t-test, $p>0.05$).

**Supplemental Table. The genotypes of flies used in each experiment.**

| | | |
|---|---|---|
| Figure 1A | control, Antimycin | elav-Gal4/+;;UAS-ATeam/+ |
| Figure 1C | elav/ATeam<br>control RNAi | elav-Gal4/+;;UAS-luciferase$^{iai16-2}$/UAS-ATeam |
| | elav/ATeam<br>milton RNAi | elav-Gal4/UAS-milton RNAi$^{v42508}$;;UAS-ATeam/+ |
| Figure 1D | 5 days, 30 days, 50 days | elav-Gal4/+;;UAS-ATeam/+ |
| Figure 2A | 5 days, 30 days, 50 days | elav-Gal4/+ |
| Figure 2B | 5 days, 30 days | elav-Gal4/+ |
| Figure 2C | 5 days, 30 days | elav-Gal4/+ |
| Figure 2D | elav/ATeam<br>control RNAi | elav-Gal4/+;;UAS-luciferase$^{B31603}$/UAS-ATeam |
| | elav/ATeam<br>Pfk RNAi | elav-Gal4/+; UAS-Pfk RNAi$^{B36782}$/+; UAS-ATeam/+ |
| Figure 2E | 5 days, 50 days | elav-Gal4/+; UAS-Pfk RNAi$^{v101887}$/+; UAS-ATeam/+ |
| Figure 2F | 5 days, 50 days | elav-Gal4/+; UAS-hGlut3/+; UAS-ATeam/+ |
| Figure 2G | elav/w1118 | elav-Gal4/+ |
| | elav/hGlut3 | elav-Gal4/+; UAS-hGlut3/+ |
| Figure 3A | 7 days, 35 days | elav-Gal4/+ |
| Figure 3B | 7 days, 35 days | elav-Gal4/+ |
| Figure 3C | elav/w1118 | elav-Gal4/+ |
| | elav/hGlut3 | elav-Gal4/+; hGlut3/+ |
| Figure4A | elav/w1118 | elav-Gal4/+ |
| | elav/hGlut3 | elav-Gal4/+; hGlut3/+ |
| Figure4B | elav/w1118 | elav-Gal4/+ |
| | elav/hGlut3 | elav-Gal4/+; hGlut3/+ |
| Figure 4C | elav/w1118 | elav-Gal4/+ |
| | elav/hGlut3 | elav-Gal4/+; hGlut3/+ |
| Figure 4D | elav/w1118 | elav-Gal4/+ |
| | elav/hGlut3 | elav-Gal4/+; hGlut3/+ |
| S1A | 5 days, 30 days | elav-Gal4/+;;UAS-ATeam/+ |
| S1B | 5 days, 30 days | elav-Gal4/+ |
| S2 | elav/ATeam | elav-Gal4/+;;UAS-ATeam/+ |
| | elav/ATeam, hGlut3 | elav-Gal4/+; UAS-hGlut3/+; UAS-ATeam/+ |
| S3A | 7 days, 35 days | elav-Gal4/+ |
| S3B | 7 days, 35 days | elav-Gal4/+ |
| S4A | Regular, 10%, 1% | elav-Gal4/+ |
| S4B | Regular, 10%, 1% | elav-Gal4/+ |
| S5 | elav/w1118 | elav-Gal4/+ |
| | elav/hGlut3 | elav-Gal4/+; hGlut3/+ |
